# Human olfactory organoids as an *in vitro* model of the olfactory epithelium

**DOI:** 10.1101/2025.10.21.679087

**Authors:** Kang-Hoon Kim, Ankit Chauhan, Michael A. Kohanski, Nithin D. Adappa, James N. Palmer, Danielle R. Reed, Noam A. Cohen, Peihua Jiang, Hong Wang, Jennifer E. Douglas

## Abstract

- A novel human olfactory epithelial organoid model has been generated from primary cell culture.
- Human olfactory organoids exhibit immunostaining and gene expression for olfactory cells.
- Olfactory organoids mobilize Ca^2+^ in response to odorants, a surrogate for neuronal function.

## Introduction

The sense of olfaction exists to perceive information about chemicals in the environment regarding safety, nutrition, and potential toxins. Perception is initiated at the periphery in the olfactory epithelium (OE), a neuroepithelium capable of regeneration throughout life. Injury of the OE can lead to olfactory dysfunction (OD), which has been linked to increased rates of depression, decreased quality of life, and increased frailty and mortality.^1,2^ The etiology of OD is varied and includes normal aging, neurodegenerative disorders, congenital conditions, head trauma, sinonasal inflammatory disease, and upper respiratory viral infection. However, an understanding of the mechanisms underlying normal olfaction and OD is lacking and has not been systematically investigated in humans due to restricted access to tissue and the lack of a human-cell-based model.

3D organoid culture of various tissue types is a demonstrated approach to studying the pathophysiology of disease.^3^ Mouse tissue^4 5^ has been a common source, but success using human tissue has been limited. Additionally, many questions about human biology and disease cannot be directly addressed using non-human models. The culture of human OE was demonstrated previously in primary cell culture;^4^ however, the resulting colonies did not demonstrate all OE cell types, OE architecture, or functionality, and did not sustain culture beyond day 14. We therefore sought to grow human OE in 3D culture as a novel *in vitro* model for studying OE biology and OD.

## Methods

### Preparation of human olfactory organoids

Institutional Review Board Approval was obtained from the University of Pennsylvania. Superior turbinate tissue was obtained from patients undergoing surgery for skull base tumors not involving nor obstructing the olfactory cleft. Inferior turbinate biopsies were obtained as a non-OE control. Objective olfactory testing was performed pre-operatively using the B-SIT (Brief Smell Identification Test^6^) to confirm normosmia. Organoids were generated by adapting previously described methods.^7,8^ Briefly, specimens were collected in DMEM/F12 medium supplemented with 1x Glutamax and 10mM HEPES before washing with 1x PBS supplemented with penicillin-streptomycin mix (Gibco, ThermoFisher Scientific, Waltham, MA). Two milliliters of 0.25% Trypsin-EDTA were added, and the specimen was homogenized and incubated at 37°C for 30 minutes. Enzymatic dissociation was arrested by the addition of DMEM/F12 medium supplemented with 10% FBS (Hyclone, Marlborough, MA) and tubes were centrifuged at 1200 rpm for 10 minutes. The supernatant was discarded, and pellet resuspended in growth medium (DMEM/F12+1x Glutamax supplemented with R-spondin (200ng/ml; R&D Systems, Minneapolis, MN), Wnt3a (50ng/ml; R&D Systems), Noggin (100ng/ml; R&D Systems), Y27632 (10µM; Sigma-Aldrich, Burlington, MA), EGF (50ng/ml; Thermofisher, Waltham, MA), bFGF (20ng/ml; Thermofisher), N2 supplement (1%; Gibco), B27 supplement (2%; Gibco), antibiotic mix (1x; Gibco), and HEPES (10mM, Gibco)). The sample was filtered and the single cell suspension mixed with Matrigel (Corning, Corning, NY) and transferred to a 24-well ultralow attachment plate (Corning). Inferior turbinate specimens were cultured following an identical protocol as a negative control.

### Immunofluorescence and RT-qPCR

Immunostaining was performed on the organoids using antibodies against OE markers to determine if the organoids contain OE cells: NCAM (immature and mature olfactory sensory neurons (OSNs)), olfactory marker protein (OMP; mature OSNs), SOX2 (globose basal cells (GBCs) and sustentacular cells (SCs)), keratin-5 (K5; horizontal basal cells (HBCs)), and keratin-8 (K8; SCs) (**Supplemental Table 1**). Nuclear staining was performed using DAPI (4′,6-diamidino-2-phenylindole) reagent (Thermofisher). RT-qPCR was performed relative to GAPDH to quantify gene expression of OE cell types: OMP (mature OSNs), SOX2 (GBCs and SCs), keratin-5 (KRT5) (HBCs), and Ezrin (SCs) (**Supplemental Table 2**).

### Live cell calcium imaging

Organoids of at least 28 days maturity were loaded with 5µM Fluo-4 AM (AAT Bioquest, Pleasanton, CA) as described previously.^9^ Imaging was performed in 1x Hanks’s Balanced Salt Solution (HBSS) buffered with 20mM HEPES and containing 1.8mM Ca^2+^. Prior to Fluo-4 AM loading, the organoids were transferred to Geltrex (Thermofisher) coated 8-well chambered glass slides (Cellvis, Mountain View, CA) and allowed to stabilize overnight. Organoids were loaded with 5 µM Fluo-4 AM for 90 minutes at 25°C and imaged with an Olympus IX-83 microscope (20x 0.75 NA Plan Apo objective) with FITC filters (Chroma technologies, Rockingham, VT) in excitation and emission filter wheels (Sutter Lambda LS), Orca Flash 4.0 sCMOS camera (Hamamatsu, Tokyo, Japan), Meta-Fluor (Molecular Devices, Sunnyvale, CA), and XCite 120 LED Boost (Excelitas Technologies, Pittsburgh, PA). Imaging was analyzed using FIJI/Image-J (National Institute of Health, Bethesda, MD). The odorant DMSO stocks were diluted in Ca^2+^-containing 1x HBSS. Organoids were sequentially exposed to vehicle (0.25% DMSO), 1mM odorant, and 100 µM ATP.

## Results

Staining of the generated olfactory organoids demonstrates the presence of OE cell-type markers: OMP (mature OSNs) and NCAM (immature and mature OSNs) (**Figure 1A-D)**, as well as SOX2 (GBCs and SCs), K5 (HBCs), and K8 (SCs) (data not shown). NCAM+ and OMP+ cells have neuron-like morphology with a dendritic knob, cell body, and long axon, typical components of an OSN. RT-qPCR demonstrates expression of markers for mature OSNs (OMP), SCs (SOX2, EZRIN), GBCs (SOX2), and HBCs (KRT5) (**Figure 1E**).

**Figure 1.**
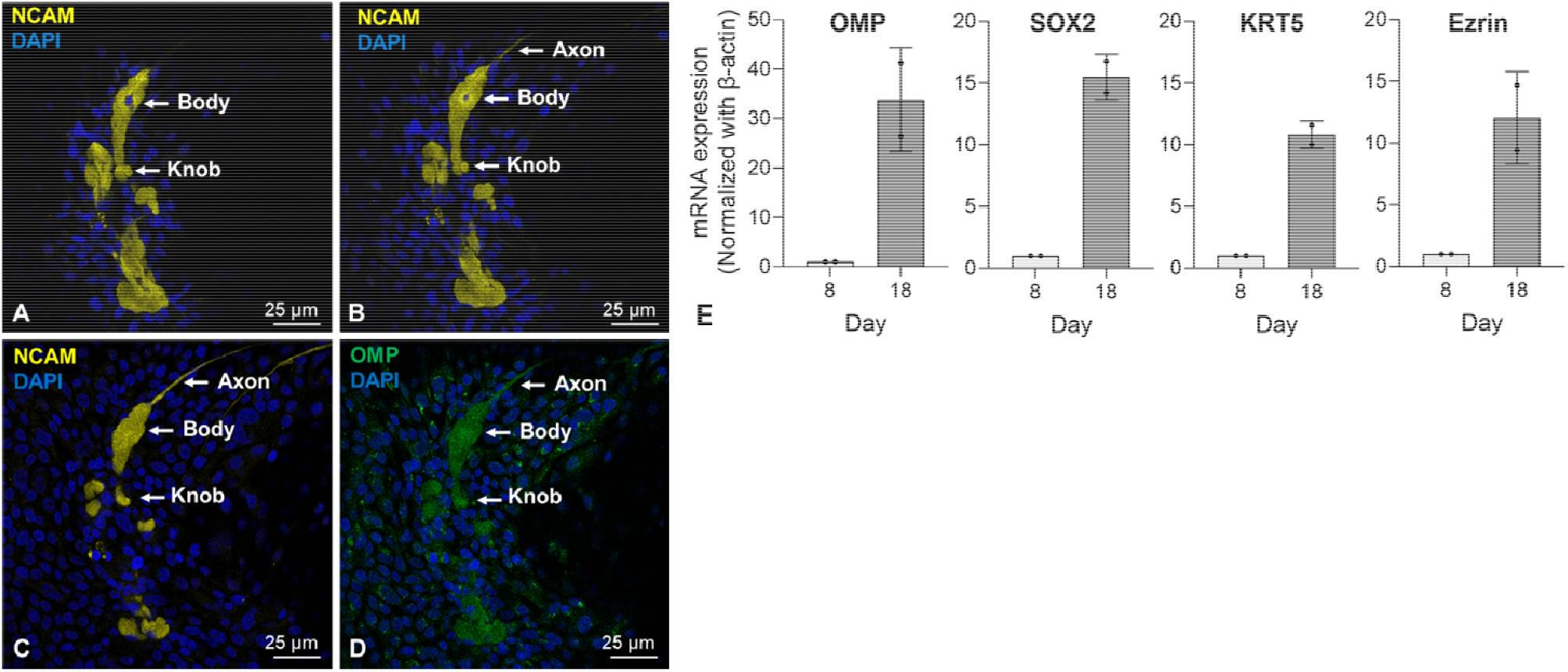
Immunostaining and RT-qPCR of human olfactory organoids demonstrate expected o factory cell types. Immunostaining of a human olfactory organoid with serial scan showing an OMP+ (**D**) and NCAM+ (**A-C**) OSN with a dendritic knob, cell body, and axon. RT-qPCR demonstrates expected cell types – OMP+ (mature OSNs), SOX2+ (GBCs and SCs), KRT5 (HBCs), and EZRIN+ (SCs) (**E**).

Live cell calcium imaging demonstrates olfactory organoids mobilize Ca^2+^ in response to odorants, suggesting organoids contain functional OSNs (**Figure 2A**). As a control, organoids cultured from inferior turbinate (non-OE) tissue do not mobilize Ca^2+^ when stimulated with odorants (**Figure 2B**).

**Figure 2.**
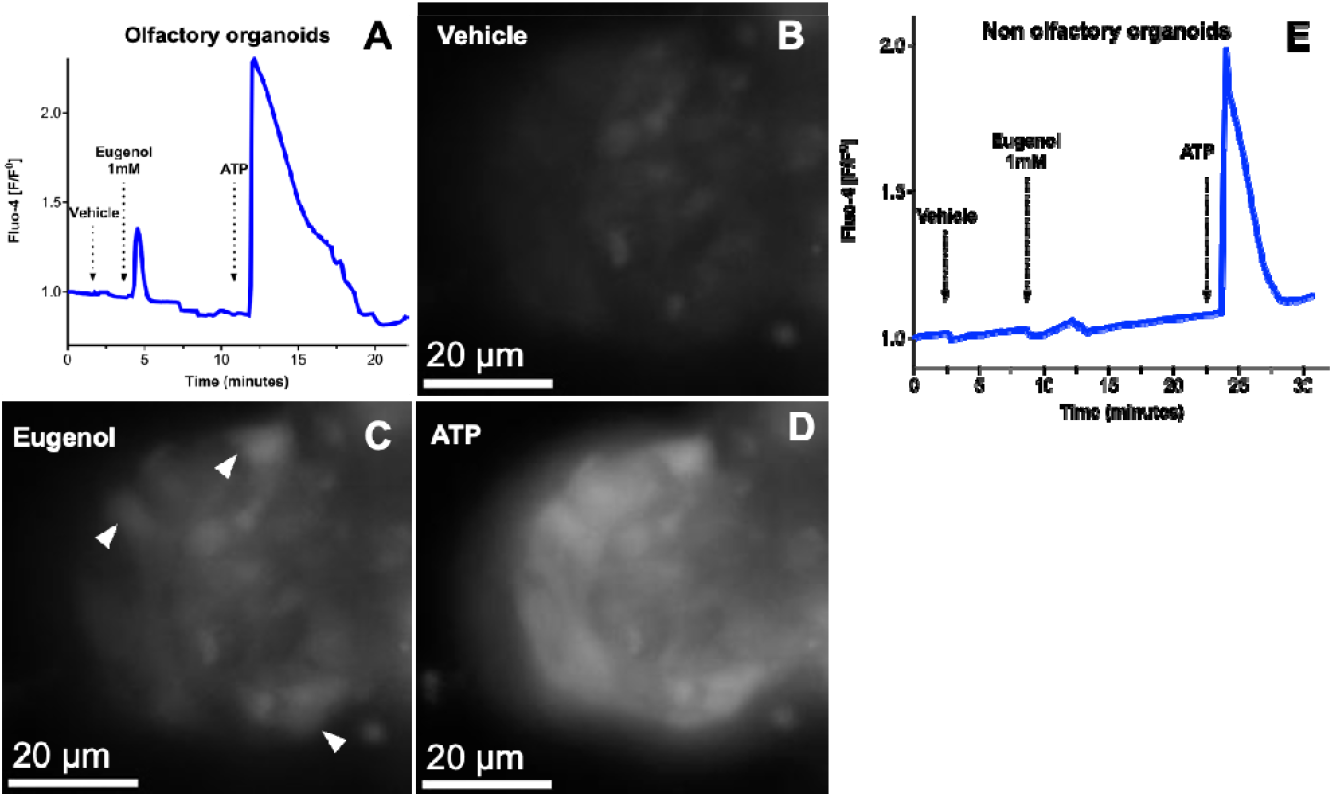
Human olfactory organoids mobilize Ca^2+^ in response to odorants. Olfactory organoids loaded with fluo-4 were imaged for Ca^2+^ response to 1mM eugenol (**A, C**). Vehicle (HBSS with 0.25% DMSO) (**B**) and 100µM ATP (**D**) were used as negative and positive controls. Non-olfactory organoids did not mobilize Ca^2+^ in response to 1mM eugenol (**E**).

## Discussion

Human superior turbinate tissue can be grown in 3D culture as a novel model for studying human OE biology. Organoids show the expected OE cell types in structure and gene expression, and mobilize Ca^2+^ in response to odorant exposure, a surrogate for function.^10^ This suggests our organoids contain functional OSNs, further supporting the model. In summary, preliminary data support a novel human olfactory organoid model, which may help investigate OE biology in normal and diseased states, with potential therapeutic insights.

## Acknowledgements

We would like to acknowledge Robert J. Lee, PhD and his laboratory for their guidance in the live cell calcium imaging detailed in the manuscript.

## Disclosure Statement

The authors have no conflicts of interest to disclose.

This project was presented at the 2025 American Rhinologic Society meeting at the American Academy of Otolaryngology – Head and Neck Surgery.

## Tables

None

## Supplemental Tables

**Table 1.**
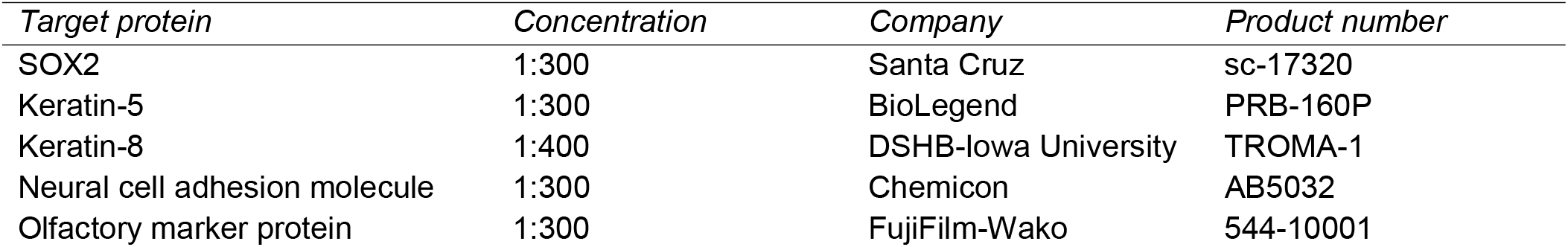
Antibodies.

**Table 2.**
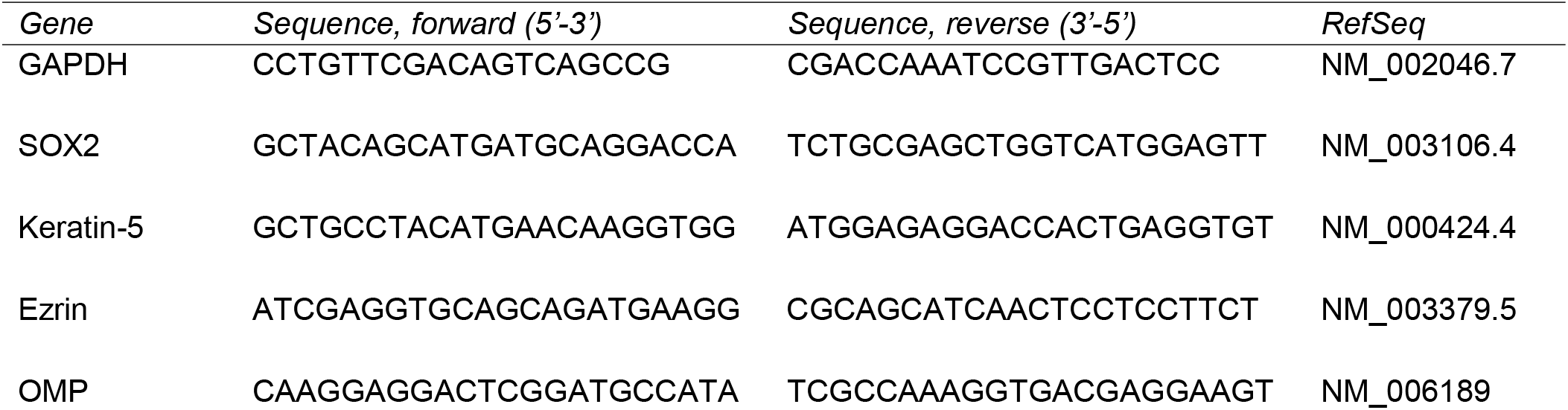
RT-qPCR primers.

