## Supplementary Table for "Human olfactory organoids as an *in vitro* model of the olfactory epithelium"

**Supplemental Tables**

| **Table 1. Antibodies** | | | |
| --- | --- | --- | --- |
| *Target protein* | *Concentration* | *Company* | *Product number* |
| SOX2 | 1:300 | Santa Cruz | sc-17320 |
| Keratin-5 | 1:300 | BioLegend | PRB-160P |
| Keratin-8 | 1:400 | DSHB-Iowa University | TROMA-1 |
| Neural cell adhesion molecule | 1:300 | Chemicon | AB5032 |
| Olfactory marker protein | 1:300 | FujiFilm-Wako | 544-10001 |

| **Table 2. RT-qPCR primers** | | | |
| --- | --- | --- | --- |
| *Gene* | *Sequence, forward (5’-3’)* | *Sequence, reverse (3’-5’)* | *RefSeq* |
| GAPDH | CCTGTTCGACAGTCAGCCG | CGACCAAATCCGTTGACTCC | NM_002046.7 |
| SOX2 | GCTACAGCATGATGCAGGACCA | TCTGCGAGCTGGTCATGGAGTT | NM_003106.4 |
| Keratin-5 | GCTGCCTACATGAACAAGGTGG | ATGGAGAGGACCACTGAGGTGT | NM_000424.4 |
| Ezrin | ATCGAGGTGCAGCAGATGAAGG | CGCAGCATCAACTCCTCCTTCT | NM_003379.5 |
| OMP | CAAGGAGGACTCGGATGCCATA | TCGCCAAAGGTGACGAGGAAGT | NM_006189 |
